# Frass fertilizers from mass-reared insects: species variation, heat treatment effects, and implications for soil application

**DOI:** 10.1101/2023.09.28.559882

**Authors:** Nadine Praeg, Thomas Klammsteiner

## Abstract

Insect farming has gained popularity as a resource-efficient and eco-friendly method of managing organic wastes by converting them into high-quality protein, fat, and frass. Insect frass is a powerful organic fertilizer, enriching the soil with essential plant nutrients and enhancing plant defense mechanisms through chitin stimulation. Given the importance of frass commercialization for many insect farmers and the use of increasingly diverse organic wastes as insect feedstock, the need for legal guidelines to enable clean production practices has emerged. The recent introduction of a legal definition for frass and heat treatment requirements by the EU commission marks a significant step towards standardizing its quality. However, frass composition is influenced by numerous factors, and little is known about the processes shaping its nutritional profiles and contributing to its maturation. Here, we analyzed the physicochemical, plant-nutritional, and microbiological properties of black soldier fly, yellow mealworm, and Jamaican field cricket frass from mass-rearing operations and assessed the impact of hygienizing heat treatment. Frass properties varied significantly across insect species, revealing concentrations of plant available nutrients reaching as high as 7000 µg NH_4_^+^, 150 µg NO_2_-NO_3_--N, and 20 mg P per g of total solids. Heat treatment affected microbial activity by reducing basal respiration and microbial biomass carbon, but also reducing viable counts of pathogenic *E. coli* and *Salmonella* sp. In terms of microbiome composition, alpha diversity showed no significant differences between fresh and heat-treated frass samples within each insect species, but significant distinctions were observed across the three insect species. The soil application of frass reactivated and boosted soil microbial activity, suggesting no long-term detrimental effects on microorganisms. These results further highlight the potential of insect frass as nutrient rich organic fertilizer, with promising benefits for soil health and nutrient cycling.

Graphical abstract:
Physicochemical and microbiological assessment of frass from black soldier fly (BSF), yellow mealworm (YMW), and Jamaican field cricket (JFC) and the impact of heat treatment. TS = total solids, VS = volatile solids, TDS = total dissolved solids, BR = basal respiration, C_mic_ = microbial biomass carbon, MQ = metabolic quotient. n.s.: p > 0.05, *: p <= 0.05, **: p <= 0.01, ***: p <= 0.001, ****: p <= 0.0001, o.o.R. = out of range, b.d.l. = below detection limit.

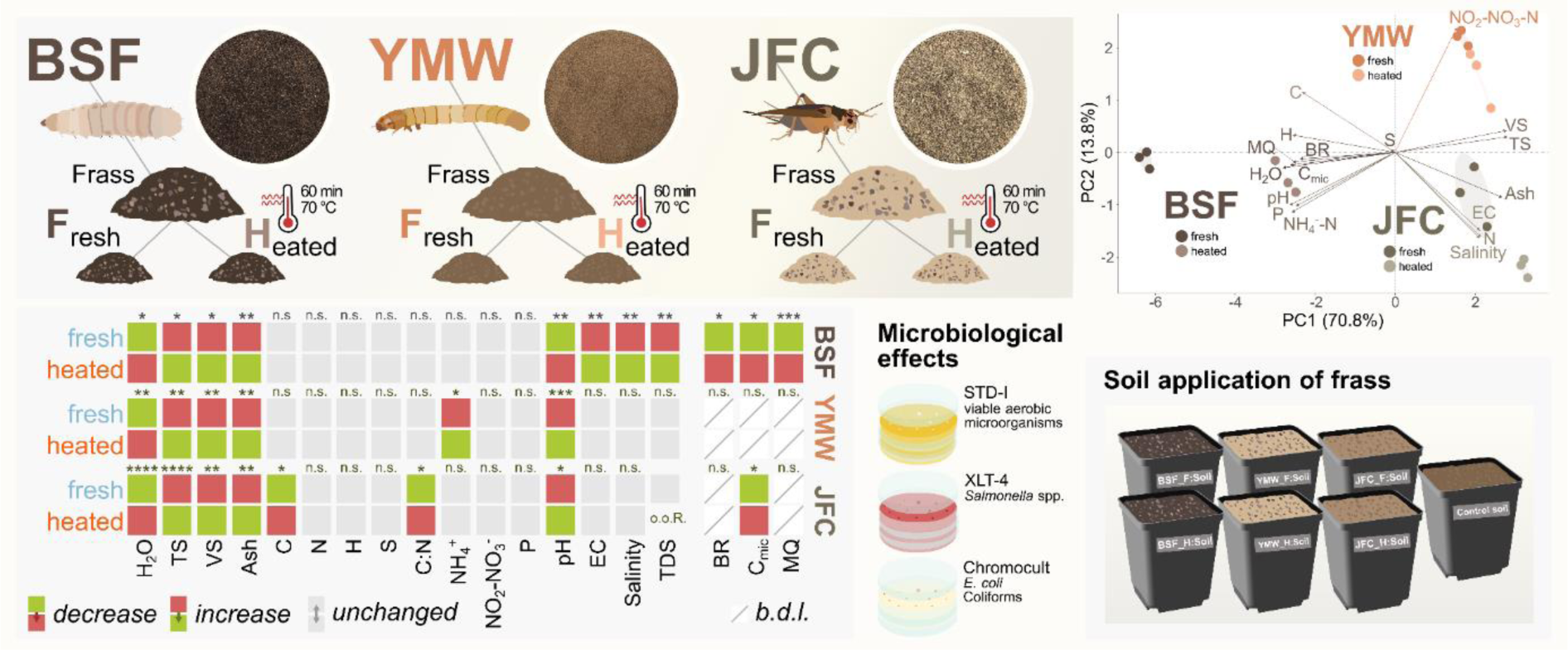

## 1. Introduction

Large-scale insect farming has emerged as a promising means to address prevalent socio-ecological problems. Compared to traditional livestock, it requires less land and water, yet achieves higher reproduction and conversion rates while generating lower greenhouse gas emissions (van Huis and Oonincx, 2017). Furthermore, it could play a pivotal role in promoting a circular economy by transforming organic waste (e.g., food wastes and manure) into valuable protein and fat, thereby minimizing resource consumption and establishing an efficient loop within the food and feed production system (Cadinu et al., 2020; Walter et al., 2020). Among the vast range of edible insects (Jongema, 2017), species such as the black soldier fly (BSF; *Hermetia illucens*, Linnaeus 1758), the yellow mealworm (YMW; *Tenebrio molitor*; Linnaeus, 1758), and Jamaican field cricket (JFC; *Gryllus assimilis*; Fabricius, 1775) have become popular among insect farmers in Western countries (Wilkie, 2018). Their popularity is primarily owed to their ability to efficiently convert organic matter and their beneficial nutritional content (Huis, 2013). While insect products for human consumption are still viewed as niche products that often evoke aversion in potential consumers, insects as feed for aquaculture, poultry, and pigs are more readily accepted (Verbeke et al., 2015).

Insect farming primarily focuses on protein and fat production, but it also inevitably generates rearing residues that may significantly contribute to the profitability of a farm (Niyonsaba et al., 2021). These residues include excrements, exuviae, undigested substrates, and dead insects and represent the main side stream of the process (Klammsteiner et al., 2020a). This so-called insect frass has been shown to have wide-ranging beneficial effects on plants and is mainly sold as organic fertilizer (Ferruzca-Campos et al., 2023; Houben et al., 2020; Menino et al., 2021). Depending on the farmed species and the substrate used to grow the insects, the general composition of frass can be highly diverse. However, its fertilizing effect is associated with the high content of organic carbon, nitrogen, and phosphorus that are comparable to other organic fertilizers (Beesigamukama et al., 2022). In addition, the chitin contained in shed skins or dead insects has been shown to improve plant immune responses and stress tolerance by simulating contact with potential pest insects (Barragán-Fonseca et al., 2022).

Microorganisms introduced into the frass, primarily via insect feces, play a crucial role in enhancing decomposition processes (Houben et al., 2020; Klammsteiner et al., 2020b). These microbes may also contribute to making the frass more similar to the gut microbiome of the insects (Gold et al., 2020). However, farmed insect species are generally selected for their rapid development, aiming to shorten rearing cycles. On average it takes approx. 20 days for BSF larvae (Heussler et al., 2022), 67 days for YMW larvae (Rumbos et al., 2021), or 60 days for JFC (Kulma et al., 2022) to reach a harvest-ready stage. Consequently, their rapid growth and constant supply of fresh feed leaves limited time for microbes and insects to effectively modify the accumulated frass, resulting in a comparatively immature compost (Beesigamukama et al., 2022). The larvae of the BSF, a commonly used insect species in waste management (Liu et al., 2022), naturally aggregate in high densities to improve feeding efficiency and are known to generate temperatures of up to 50 °C when reared at large scale (Shishkov et al., 2019; Ushakova et al., 2018). Although such elevated temperatures may even induce stress responses in insects, they are not comparable to the temperature profiles of traditional composting methods and are therefore insufficient for the microbiological stabilization of the frass (Insam et al., 2023). The composting process typically induces one or more temperature peaks and drastically increases temperatures within compost heaps to 70 °C and higher, thus, naturally hygienizing the composted material (Zhou et al., 2022). This temperature rise ensures the elimination of potentially harmful pathogens and weed seeds, promoting the microbiological maturation of the compost and producing a stable end product.

To increase product safety and reduce potential health hazards from pathogens in insect products, the EU commission has established a detailed definition for insect frass, categorizing it in the same group with processed animal manure. This regulation introduces first hygienic standards for insect frass, requiring farmers to heat-treat frass at temperatures of 70 °C for at least 60 min (Regulation (EU) 2021/1925, 2021). Although this mandatory pretreatment should ensure pathogen removal, it may also inhibit microbial activity beneficial to the frass’ nutrient content and alter its value as soil fertilizer. So far, only one study has investigated the effect of heat treatment on the frass of black soldier fly larvae and found that it was successful in reducing Enterobacteriaceae, *Salmonella*, and *Clostridium perfringens* below the detection limit (Van Looveren et al., 2022). However, total viable counts only decreased by 1-log and bacterial endospores were unaffected. With the rapid expansion of the insect farming sector and the value of commercializing rearing residues, a thorough assessment of risks and opportunities becomes imperative.

In this study, we conducted a comprehensive characterization and comparison of physicochemical and microbiological features of frass from three widely farmed insect species (BSF, YMW, JFC). To explore the effects of hygienization, we subjected the frass to heat treatment following legislative guidelines. By using 16S rRNA gene amplicon sequencing, we evaluated differences in bacterial communities between insect species and before and after heat treatment. Furthermore, we assessed how this hygienization process influences the frass’s suitability as an organic soil amendment by conducting a soil incubation trial and screening the frass-amended soils thereafter.

## 2. Materials and Methods

### 2.1. Origin and pretreatment of the frass

Fresh, untreated frass from BSF (Figure 1A), YMW (Figure 1B), and JFC (Figure 1C) was obtained from commercial insect farmers in Austria (BSF and YMW) and Croatia (JFC). Upon arrival, the frass was stored at −20 °C and, prior to its use, gently thawed over 24 h at 4 °C. For the heat treatment, the frass was evenly spread out on large glass petri dishes at a layer height of approx. 1 cm and incubated at 70 °C for 1 h in a preheated drying cabinet (Memmert, Schwabach, Germany). Following this procedure, the heat-treated frass was transferred into paper bags and left to cool down to room temperature before further use.

**Figure 1.**
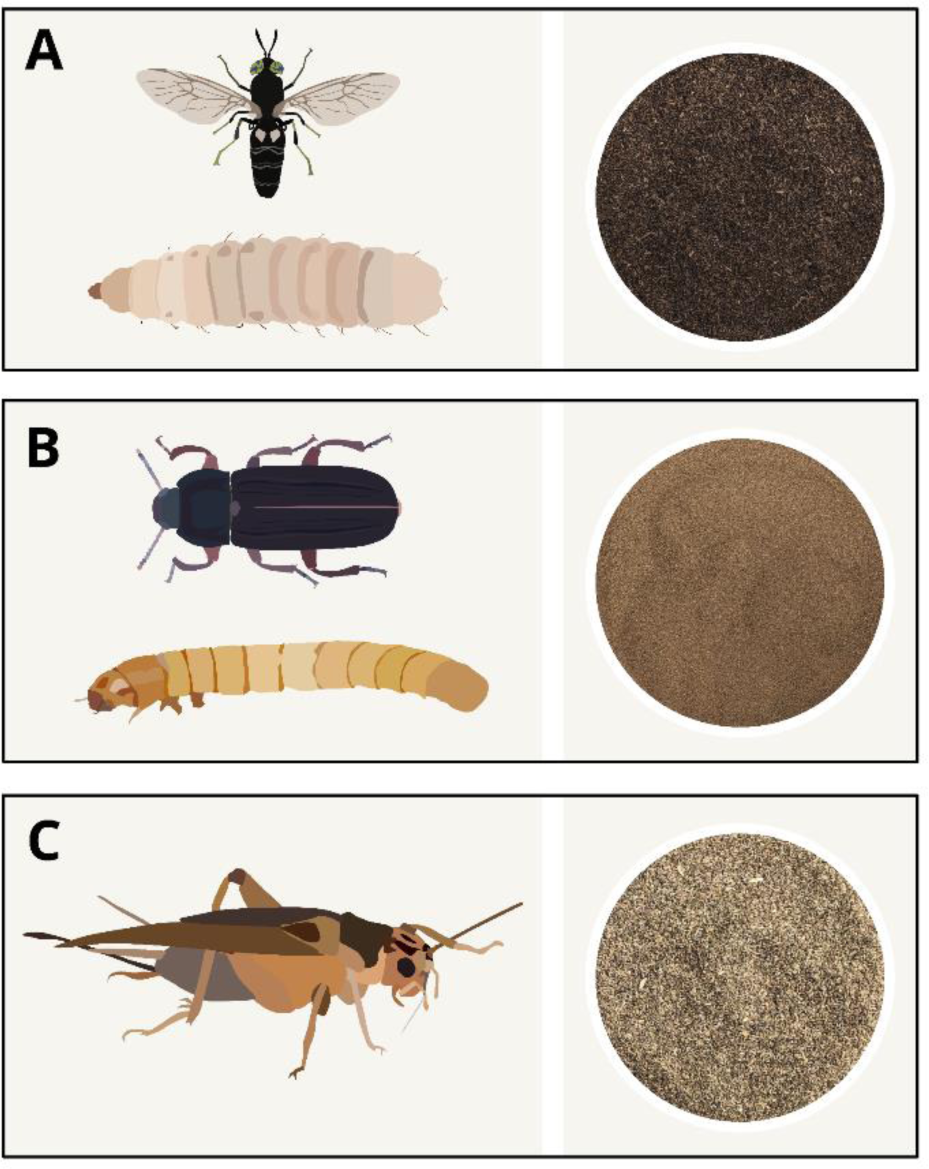
Three insect species that are approved in the EU for food and feed applications, and also serve as frass producers. **A.** Black soldier fly (*Hermetia illucens*; Linnaeus, 1758) adult, larva, and its frass **B.** Yellow mealworm (*Tenebrio Molitor*; Linnaeus, 1758) adult, larva, and its frass **C.** Jamaican field cricket (*Gryllus assimilis*; Fabricius, 1775) adult and its frass. The photos of frass represent the actual material used in this study.

### 2.2. Physicochemical parameters

#### 2.2.1. Water, total solids, volatile solids, and ash content

Water and total solids (TS) content were determined gravimetrically by calculating the loss in mass before and after drying the samples (n = 3) at 105 °C for 24 h in a drying cabinet (UN110, Memmert, Schwabach, Germany). To determine the volatile solids (VS) and ash content, the TS fraction was finely ground using a pestle and mortar and incinerated in a muffle furnace (CWF 1000, Carbolite, Neuhausen, Germany) at 550 °C for 5 h (n = 3). The loss in mass was interpreted as VS fraction, while the residues were considered as the ash content.

#### 2.2.2. pH, electrical conductivity, and salinity

Samples (n = 4) were weighted into 50 mL plastic falcon tubes, mixed with a. deion. in a ratio of 1:12.5 (w/v), and incubated at room temperature overnight before measurement. A 774 pH Meter (Metrohm, Herisau, Switzerland) was used to measure pH in the diluted samples. Electrical conductivity (EC), salinity, and total dissolved solids (TDS) were measured in the same samples using a LF330/SET conductivity electrode (WTW, Weilheim in Oberbayern, Germany).

#### 2.2.3. Elemental analysis (CHNS)

Part of the dried biomass resulting from TS determination was ground and sent to the Department of Waste and Resource Management (TU Wien, Vienna, Austria) for elemental analysis. The CHNS content was determined using a Vario MACRO elemental analyzer (Elementar, Langenselbold, Germany). Ca. 15 mg of sample material wrapped in a tin capsule was combusted at 1150 °C and the resulting combustion gas was separated through an adsorption column reducing NOx to Cu and subsequently N2. Sequential desorption was induced by heating the absorption column and gasses were measured by a thermal conductivity detector. He 5.0 was used as a carrier gas.

#### 2.2.4. Plant-available ammonium, nitrate and phosphorus content

Plant-available ammonium (NH_4_^+^-N; µg g^-1^ TS), nitrate (NO_2_-NO_3-_-N; µg g^-1^ TS) were determined based on a modified Berthelot and cadmium reduction method, respectively, after shaking 2 g frass or 7.5 g soil:frass mixture in 30 mL KCl [1 M] for 1 h at 120 rpm. Plant-available phosphorus (ortho-phosphate, µg g^-1^ TS) was determined by applying the Olsen method and shaking 0.4 g frass or 2 g soil:frass mixture in 40 ML LiCl [0.4 M] for 16 h at 150 rpm. All extracts from frass and soil:frass mixtures were filtered (Macherey & Nagel 615¼, 150 mm filter paper) and analyzed for the respective nutrient content with a Continuous Flow Analyzer (CFA, Skalar, Netherlands).

### 2.3. Microbiological parameters

#### 2.3.1. Basal respiration and microbial biomass

Basic soil respiration and substrate-induced respiration (SIR) (for the calculation of microbial biomass) were determined on an EGA61-Soil respiration Device (ADC BioScientific, UK). Frass and soil:frass mixtures were filled into acrylic glass tubes, closed with polystyrene foam pads and aerated with a continuous stream of ambient air (humidified and tempered to 22 °C). The CO_2_ released from the samples was recorded for 6 h with an infrared gas analyzer (IRGA) to calculate the basic soil respiration (BR [µg CO2 g^-1^ TS h^-1^]). Afterwards, glucose (1%, w/w dry weight) was added to the samples, and the CO_2_ release was further recorded for 12 h (substrate-induced respiration method). The maximum CO_2_ release was used to calculate the microbial biomass (C_mic_ [µg C g^-1^ TS]) according to Anderson and Domsch (1978). The metabolic quotient (MQ) was calculated as the quotient of basic soil respiration (BR) and microbial biomass (C_mic_).

#### 2.3.2. Microbial counts

Standard I nutrient agar was prepared from 15 g peptone, 3 g yeast extract, 6 g NaCl, 1 g glucose, 12 g agar, and adjusted to a pH of 7.5 ± 0.2 using HCl [0.5 M] to determine the total plate count of viable aerobic bacteria. Chromocult® TBX (Tryptone Bile X-glucuronide) agar (Merck, Darmstadt, Germany) and XLT4 agar (Merck, Darmstadt, Germany) prepared according to the manufacturer’s instructions were used to detect and quantify *E. coli* and *Salmonella* sp. For the dilution series, 1 g sample biomass was sequentially diluted in sterile Ringer’s solution (Merck, Darmstadt, Germany) up to a dilution level of 10^-8^. After selecting three appropriate levels of dilution for each type of medium, 50 µL of the resulting dilutions were plated onto the respective agar plates. All plates were incubated at 37 °C and inspected after 24 and 48 h for the quantification of CFUs and detection of pathogens.

#### 2.3.3. DNA extraction and marker gene sequencing for bacterial and fungal communities

DNA was extracted from 150 (BSF), 200 (YMW), and 300 mg (JFC) of both fresh and treated frass (n = 3) using the NucleoSpin Soil Kit (Macherey-Nagel, Düren, Germany) and following the manufacturer’s protocol. Lysis buffer SL2 was used for the cell lysis step and washed extracts were eluted in MN elution buffer. DNA yield and purity was assessed via UV-vis spectrophotometry using a NanoDrop 2000c device (Thermo Fisher Scientific, Waltham, MA, USA). All samples passing quality control were sent for 16S rRNA gene amplicon sequencing. Sequencing was carried out on the NovaSeq6000 platform (Illumina, San Diego, CA, USA) following a 2×250 bp approach and using the primer pair 515f (5′-GTGCCAGCMGCCGCGGTAA-3′) and 806r (5′-GGACTACHVGGGTWTCTAAT-3′) to target the V4 region on the 16S rRNA gene. Raw reads were processed using dada2 v.1.26.0 (Callahan et al., 2016) and classified into amplicon sequence variants (ASVs) following the latest standard operating procedure (https://benjjneb.github.io/dada2/tutorial.html). Briefly, adapter- and primer-free reads were truncated at a length of 200 bp based on the inspected quality profiles, and any reads containing Ns were discarded. After learning the error rates and sample inference using the default settings, paired-end reads were merged to construct the sequence table. Chimeras were removed using the removeBimeraDenovo() command by applying the “consensus” method. Taxonomy was assigned to the ASVs based on the SILVA trainset version 138.1 (Quast et al., 2013). ASVs consisting of less than three reads and not detected in at least 10% of the samples were removed from the data.

### 2.4. Soil incubation trial

Soil was collected from nearby agricultural land (47°15’47.6“N 11°20’24.0”E) and stored at 4 °C overnight. To remove any plant residues and stones, the soil was passed through a 4 mm sieve and its physicochemical properties were determined (Table 1).

**Table 1.** Physicochemical properties of the soil used to prepare the soil:frass mixtures. Values indicate mean ± standard deviation (n = 3).

| Measured parameter | Values |
| --- | --- |
| Water content [%] | 24.4 ± 0.2 |
| Total solids [%] | 75.6 ± 0.2 |
| Volatile solids [%] | 6.0 ± 0.1 |
| Ash [%] | 69.7 ± 0.2 |
| C [%] | 4.8 ± 0.3 |
| N [%] | 0.3 ± 0.0 |
| C:N ratio | 15.9 ± 2.1 |
| pH | 7.17 ± 0.11 |
| EC [μS $\text{cm}^{-1}$] | 76 ± 4 |
| Salinity | 0 ± 0 |
| TDS [$\text{mg L}^{-1}$] | 30.3 ± 2.0 |

The soil:frass mixtures were prepared following the recommended dosage for each type of fresh frass. For the heat-treated frass, the amounts were adjusted to match the total solids content of the untreated frass, as indicated in Table 2. For each replicate (n = 4), 200 g of sieved soil were thoroughly mixed with frass in the recommended ratio in a plastic bucket. The mixtures were transferred into nursery pots for plants (Ø_top_ = 90 mm, Ø_bottom_ = 60 mm, h = 80 mm), evenly moistened with deionized water using a spray bottle and loosely covered with cling foil. Incubation of the pots was conducted in a shaded greenhouse for 14 days with conditions set to 20 °C and 70% relative humidity. The loss of water due to evaporation was monitored continuously by weighing the pots on a portable scale (KF6000A, G&G, Kaarst, Germany) and adjusting the water content by spraying the soil surface of the pots with a. deion. (Figure S1).

**Table 2.** Mixing ratios of fresh, untreated and heat-treated frass from industrially farmed black soldier fly (BSF), yellow mealworm (YMW), and Jamaican field cricket (JFC). The dosage for heat-treated frass was adapted based on the content of total solids in fresh frass.

|  | BSF frass |  | YMW frass |  | JFC frass |  |
| --- | --- | --- | --- | --- | --- | --- |
|  | Fresh | Heated | Fresh | Heated | Fresh | Heated |
| Recommended dosage [%] | 2-3 |  | 10-15 |  | 10-15 |  |
| Total solids [%] | 57.4 ± 3.3 | 69.5 ± 1.0 | 87.9 ± 0.1 | 91.6 ± 0.4 | 85.4 ± 0.2 | 91.9 ± 0.2 |
| Frass per volume soil [%] | 2.7 | 2.2 | 12.5 | 12 | 12.5 | 11.6 |

### 2.5. Statistics and data analysis

All statistical calculations and visualizations were carried out in R v.4.1.2 (R Core Team, 2021). To assess differences in the physicochemical composition of sample groups, analysis of variance (ANOVA) was calculated using the aov() function from the R ‘*stats*’ package. For overall pairwise comparisons, Tukey’s Honestly Significant Difference (TukeyHSD) posthoc test was calculated using the glht() function in the ‘*multcomp*’ package (Hothorn et al., 2008), and summaries for statistical similarities and differences were generated using the multcompletters4() function in the ‘*multcompView*’ package (Graves et al., 2019). Principal component analysis (PCA) was performed on the normalized data of physicochemical parameters (TS, H_2_O, VS, ash, pH, EC, salinity, C, H, N, S, NH_4_^+^-N, NO_2_-NO_3-_-N, P, BR, MQ, and C_mic_) using the prcomp() function in the R ‘*stats’* package. The PCA results were visualized using the fviz_pca_biplot() function in the R ‘*factoextra*’ package (Kassambara and Mundt, 2020). Differences in alpha diversity were calculated via ANOVA and pairwise Wilcoxon Rank Sum Test using the pairwise.wilcox.test() function with Bonferroni correction from the ‘*stats*’ package. Beta diversity was calculated and visualized via principal coordinates analysis (PCoA) using the amp_ordinate() function in ‘*ampvis2*’ (Andersen et al., 2018). Linear discriminant analysis of effect size (LEfSe) for the identification of biomarkers in microbial community data was calculated using the run_lefse() function from the ‘*microbiomeMarker’* package (Yang, 2021). To test differences between groups of samples, permutational multivariate analysis of variance (PERMANOVA) based on Bray-Curtis dissimilarity matrices was calculated using the adonis2() function in the ‘*vegan’* package (Oksanen et al., 2020).

## 3. Results

The aim of this study was to characterize the physicochemical and microbiological properties of frass samples obtained from three industrially exploited insect species (Figure 1), and to emphasize the key points of differentiation among them. Given the increasing concerns about the safety of utilizing untreated rearing residues from insect farming as agricultural fertilizer, we further examined the effects of heat treatment at 70 °C for 1 h on the physicochemical and microbiological characteristics of the frass. To achieve these results, we employed a combination of chemical nutrient analyses, microbiological cultivation techniques, and biomolecular sequencing methods. In addition, we conducted a two-week incubation trial in a greenhouse to assess how the supplementation of fresh and heat-treated frass affects nutrient content and microbial activity in soil.

### 3.1. Physicochemical characterization of fresh and heat-treated frass

The frass samples significantly varied in their general composition, with the primary differentiating factor being the insect species of origin, as highlighted by the first two principal components of the PCA that explained 85% of the variance in the data (Figure 2A). The heat treatment had a comparably minor impact on the distribution of the samples, as they showed minimal divergence from their corresponding fresh counterparts upon pairwise comparison of the respective groups (Figure 2B). A detailed analysis of the physicochemical drivers is shown in Table 3. The fresh frass samples represented the condition in which the untreated material is sold by the producers (Figure 1A-C); however, they significantly varied in their water content. While BSF frass was comparatively humid with a water content of 43%, the other two types of frass ranged between 12 and 15%. The heat treatment significantly reduced the water content in BSF frass by 12%, while YMW and JFC lost approx. 4-7%. Accordingly, BSF frass exhibited the lowest total solids (TS) content at 57%, resulting in a 28% and 31% reduction compared to JFC and YMW, respectively. Comparable patterns applied to the relationship between volatile solids (VS) and ash content.

**Figure 2.**
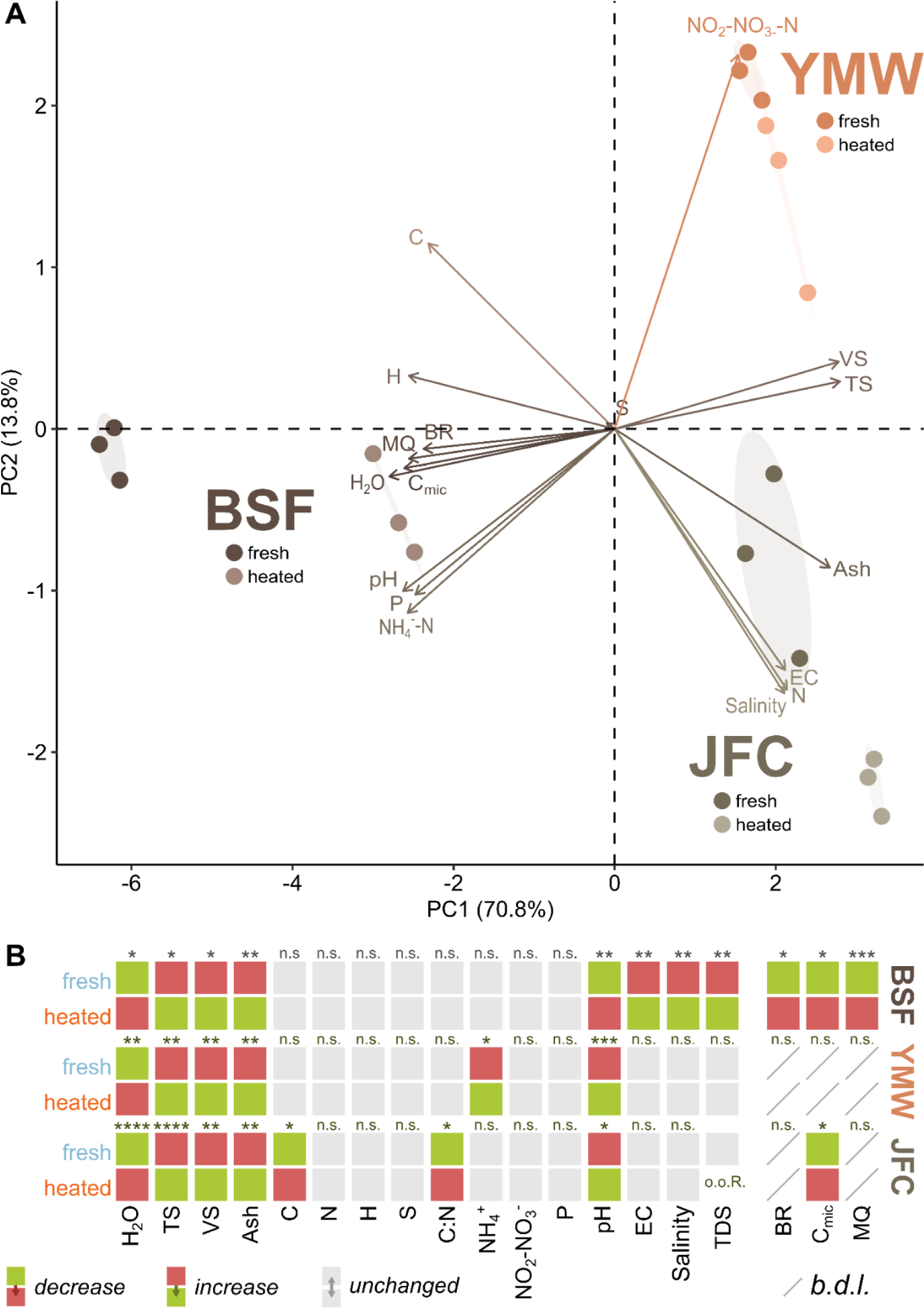
**A.** Principal component analysis of physicochemical and microbiological parameters measured in fresh and heat-treated frass samples of black soldier fly (BSF), yellow mealworm (YMW), and Jamaican field cricket (JFC) (n = 3). **B.** Heatmap showing results of pairwise t-tests for fresh and heat-treated frass samples. TS = total solids, VS = volatile solids, TDS = total dissolved solids, BR = basal respiration, C_mic_ = microbial biomass carbon, MQ = metabolic quotient. n.s.: p > 0.05, *: p <= 0.05, **: p <= 0.01, ***: p <= 0.001, ****: p <= 0.0001, o.o.R. = out of range, b.d.l. = below detection limit.

**Table 3.** Physicochemical parameters (mean ± standard deviation) before and after treating the frass at 70 °C for 1 h. Statistical differences were calculated via analysis of variance (ANOVA) followed by Tukey HSD posthoc tests for pairwise comparison of sample groups (n = 4). o.o.R. = out of range, n.s. = not significant.

|  | BSF frass |  | YMW frass |  | JFC frass |  | F-value |
| --- | --- | --- | --- | --- | --- | --- | --- |
|  | Fresh | Heated | Fresh | Heated | Fresh | Heated |  |
| Water content [%] | 42.6 ± 3.3<sup>a</sup> | 30.5 ± 1<sup>b</sup> | 12.1 ± 0.1<sup>cd</sup> | 8.4 ± 0.4<sup>de</sup> | 14.6 ± 0.2<sup>c</sup> | 8.1 ± 0.2<sup>e</sup> | $F=292.322^{***}$ |
| Total solids [%] | 57.4 ± 3.3<sup>e</sup> | 69.5 ± 1<sup>d</sup> | 87.9 ± 0.1<sup>bc</sup> | 91.6 ± 0.4<sup>ab</sup> | 85.4 ± 0.2<sup>c</sup> | 91.9 ± 0.2<sup>a</sup> | $F=292.322^{***}$ |
| Volatile solids [%] | 50.4 ± 3.1<sup>d</sup> | 61.2 ± 0.9<sup>c</sup> | 78.7 ± 0.1<sup>ab</sup> | 82 ± 0.4<sup>a</sup> | 75.6 ± 0.1<sup>b</sup> | 80.8 ± 0.4<sup>a</sup> | $F=281.043^{***}$ |
| Ash [%] | 7.0 ± 0.3<sup>e</sup> | 8.4 ± 0.1<sup>d</sup> | 9.2 ± 0<sup>c</sup> | 9.6 ± 0<sup>bc</sup> | 9.9 ± 0.2<sup>b</sup> | 11.1 ± 0.2<sup>a</sup> | $F=223.27^{***}$ |
| C [%] | 42.8 ± 0.3<sup>a</sup> | 42.8 ± 0.3<sup>a</sup> | 42.1 ± 0.1<sup>b</sup> | 42.0 ± 0.1<sup>b</sup> | 41.3 ± 0.2<sup>c</sup> | 40.8 ± 0.3<sup>d</sup> | $F=73.365^{***}$ |
| H [%] | 5.90 ± 0.05<sup>a</sup> | 5.83 ± 0.02<sup>a</sup> | 5.55 ± 0.07<sup>bc</sup> | 5.60 ± 0.03<sup>b</sup> | 5.50 ± 0.07<sup>cd</sup> | 5.45 ± 0.03<sup>d</sup> | $F=66.872^{***}$ |
| N [%] | 2.84 ± 0.05<sup>d</sup> | 2.80 ± 0.14<sup>d</sup> | 3.84 ± 0.03<sup>c</sup> | 3.85 ± 0.03<sup>c</sup> | 6.07 ± 0.15<sup>b</sup> | 6.41 ± 0.20<sup>a</sup> | $F=853.835^{***}$ |
| S [%] | 0.92 ± 0.17 | 0.77 ± 0.01 | 0.92 ± 0.17 | 0.78 ± 0.01 | 0.98 ± 0.18 | 0.88 ± 0.01 | n.s. |
| C:N ratio | 15.1 ± 0.4<sup>a</sup> | 15.3 ± 0.8<sup>a</sup> | 11.0 ± 0.1<sup>b</sup> | 10.9 ± 0.1<sup>b</sup> | 6.8 ± 0.2<sup>c</sup> | 6.4 ± 0.2<sup>c</sup> | $F=462.91^{***}$ |
| NH<sub>4</sub><sup>+</sup>-N [μg g<sup>-1</sup> TS] | 6989 ± 372<sup>a</sup> | 5877 ± 90<sup>b</sup> | 848 ± 8<sup>c</sup> | 916 ± 16<sup>c</sup> | 2599 ± 44<sup>d</sup> | 2549 ± 41<sup>d</sup> | $F=791.633^{***}$ |
| NO<sub>2</sub>-NO<sub>3</sub>-N [μg g<sup>-1</sup> TS] | 2.95 ± 0.95<sup>c</sup> | 2.61 ± 0.40<sup>c</sup> | 153.28 ± 13.23<sup>a</sup> | 145.74 ± 7.55<sup>a</sup> | 51.74 ± 24.12<sup>b</sup> | 18.83 ± 1.97<sup>c</sup> | $F=106.992^{***}$ |
| P<sub>plant available</sub> [mg g<sup>-1</sup> TS] | 20.34 ± 1.56<sup>a</sup> | 20.86 ± 0.55<sup>a</sup> | 10.16 ± 0.14<sup>c</sup> | 10.95 ± 0.30<sup>bc</sup> | 12.73 ± 1.20<sup>b</sup> | 12.91 ± 0.98<sup>b</sup> | $F=76.587^{***}$ |
| pH | 7.66 ± 0.07<sup>a</sup> | 7.34 ± 0.01<sup>b</sup> | 6.24 ± 0.02<sup>e</sup> | 6.39 ± 0.03<sup>d</sup> | 6.57 ± 0.02<sup>c</sup> | 6.62 ± 0.01<sup>c</sup> | $F=740.976^{***}$ |
| EC [mS cm<sup>-1</sup>] | 4.13 ± 0.11<sup>d</sup> | 4.55 ± 0.06<sup>bc</sup> | 4.43 ± 0.04<sup>c</sup> | 4.66 ± 0.16<sup>bc</sup> | 4.82 ± 0.26<sup>a</sup> | 5.09 ± 0.01<sup>ab</sup> | $F=12.026^{***}$ |
| Salinity | 2.1 ± 0.08<sup>d</sup> | 2.38 ± 0.05<sup>bc</sup> | 2.3 ± 0<sup>c</sup> | 2.43 ± 0.13<sup>bc</sup> | 2.5 ± 0.14<sup>b</sup> | 2.7 ± 0<sup>a</sup> | $F=10.989^{***}$ |
| TDS [mg L<sup>-1</sup>] | 1647 ± 42<sup>c</sup> | 1816 ± 24<sup>ab</sup> | 1767 ± 16<sup>b</sup> | 1863 ± 66<sup>ab</sup> | 1881 ± 57<sup>a</sup> | o.o.R. | $F=16.99^{***}$ |

The elemental analysis revealed no significant differences in the relative content of C, H, and S among the three types of frass both before and after heat treatment. However, decisive differences in the N content were observed, with JFC frass containing up to twice as much N compared to the others, significantly shifting its C:N ratio to 6-7 as opposed to ca. 15 in BSF and 11 in YMW frass.

Fresh BSF frass demonstrated the highest NH_4_^+^-N content, reaching nearly 7000 µg g^-1^ TS. Notably, it was also the only type of frass, where the NH_4_^+^-N content was significantly reduced following heat treatment. YMW samples stood out with levels of NO_2_-NO_3_--N that were 3-7 times higher than in JFC samples and 50-55 times higher than in BSF, thereby contributing to the distinctiveness of this particular type of frass. Generally, NO_2_-NO_3_--N concentrations were not affected by heat treatment, except for JFC samples where it led to a 2.7-fold decrease from an average of 52 to 19 µg g^-1^ TS (Table 3). With more than 20 mg g^-1^ TS, plant-available P in BSF frass samples exceeded the concentrations of the other two types of frass by far; furthermore, the plant-available P content remained unaffected by heat treatment across all samples.

The fresh frass pH ranged between 6.2 and 7.7, and after heat treatment, it slightly decreased in BSF, slightly increased in YMW, and remained unchanged in JFC frass. Electrical conductivity (EC) slightly increased in all samples after heat treatment, but remained between 4.6 and 5.1. As measurements of salinity and total dissolved solids are functions of EC, they followed the same patterns.

### 3.2. Microbiological characterization of fresh and heat-treated frass

Notable microbial activity was mainly observed in fresh BSF frass, and the application of heat treatment significantly reduced this activity (Table 4). In these samples, BR declined by a factor of 23 after heating and reduced the C_mic_ to a third. In turn, the MQ as a function of C_mic_ and BR dropped from ca. 30 to 5 µg CO_2_-C h^-1^ per µg^-1^ C_mic_ after the treatment. Although significantly lower amounts of C_mic_ were quantified in fresh JFC frass, no BR was measured in these samples. Consequently, no microbial activity, as indicated by the MQ, could be observed. Microbial activity was detected in neither fresh nor heat-treated frass of YMW.

**Table 4.**
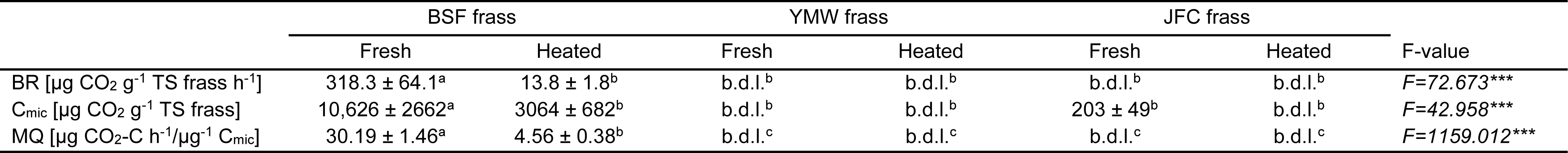
Microbiological parameters (mean ± standard deviation) before and after treating the frass at 70 °C for 1 h. Statistical differences were calculated via analysis of variance (ANOVA) followed by Tukey HSD posthoc tests for pairwise comparison of sample groups (n = 4). b.d.l. = below detection limit.

Only in BSF frass, counts of colony forming units (CFUs) of aerobically cultivable bacteria were significantly reduced from 1.3 10^9^ to 3.8 10^8^ after heat treatment (Figure 3, Supplementary Table 1). No meaningful reduction was observed in YMW and JFC frass. CFUs of *E. coli* were found in neither fresh nor heat-treated samples from any of the three insect species. However, *Salmonella* sp. was detected in fresh JFC frass, with comparably low CFU counts of 1.7 10^3^, which were reduced to beneath the detection limit after heat treatment.

**Figure 3.**
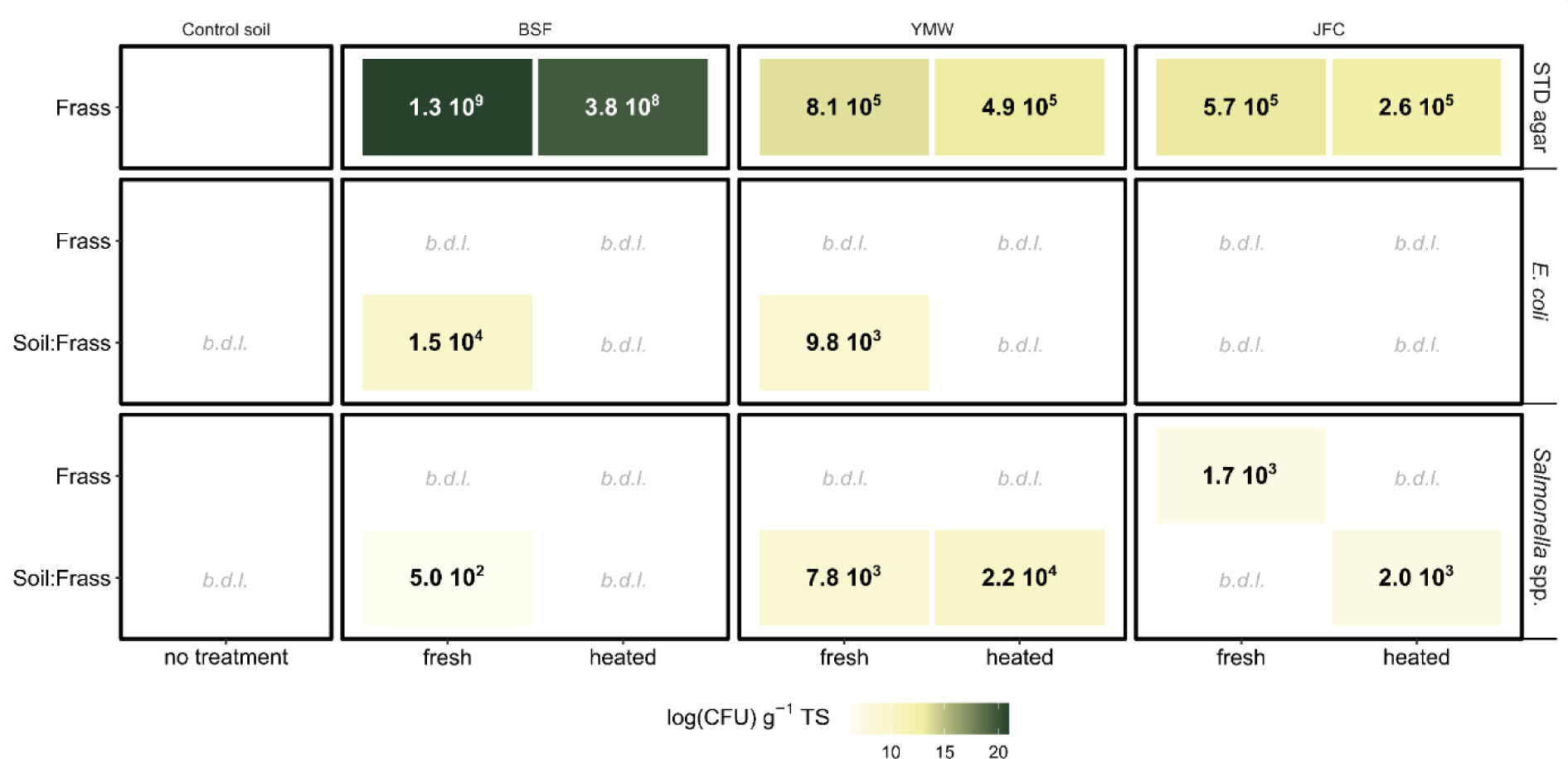
Average counts of colony forming units (CFUs) g^-1^ TS frass and soil:frass on Standard I (STD), TBX ChromoCult™ (for the detection of *E.coli*), and XLT4 (for the detection of *Salmonella* spp.) medium after 48 h of incubation (n = 3) under fresh and heated conditions. Counts have been log-transformed for the gradient fill scale. Total viable counts of aerobic microorganisms (STD agar) were only assessed for frass samples. b.d.l. = below detection limit.

### 3.3. Analysis of frass microbial communities

An average of 137,110 ± 4633 raw reads were obtained from amplicon sequencing of frass DNA extracts. After sequence preprocessing, the read count was reduced to 125,474 ± 6322 merged reads per sample for subsequent data analysis. Alpha diversity, as measured by observed species (Figure 4A) and the Shannon-Wiener index (Figure 4B), showed no significant differences between fresh and heated frass samples within each insect species, as confirmed by the Wilcoxon rank sum test. However, the frass samples, whether fresh or heated, significantly differed across the three insect species in terms of observed species (fresh: *F_2,6_* = 31.9, p < 0.001; heated: *F_2,6_* = 118, p < 0.001) and Shannon-Wiener diversity index (fresh: *F_2,6_* = 43.7, p < 0.001; heated: *F_2,6_* = 90.1, p < 0.001). The TukeyHSD test demonstrated a significant difference (p < 0.05) between JFC and BSF only after heating, but not under fresh conditions. This distinction was evident in both the observed species and the Shannon-Wiener diversity index.

**Figure 4.**
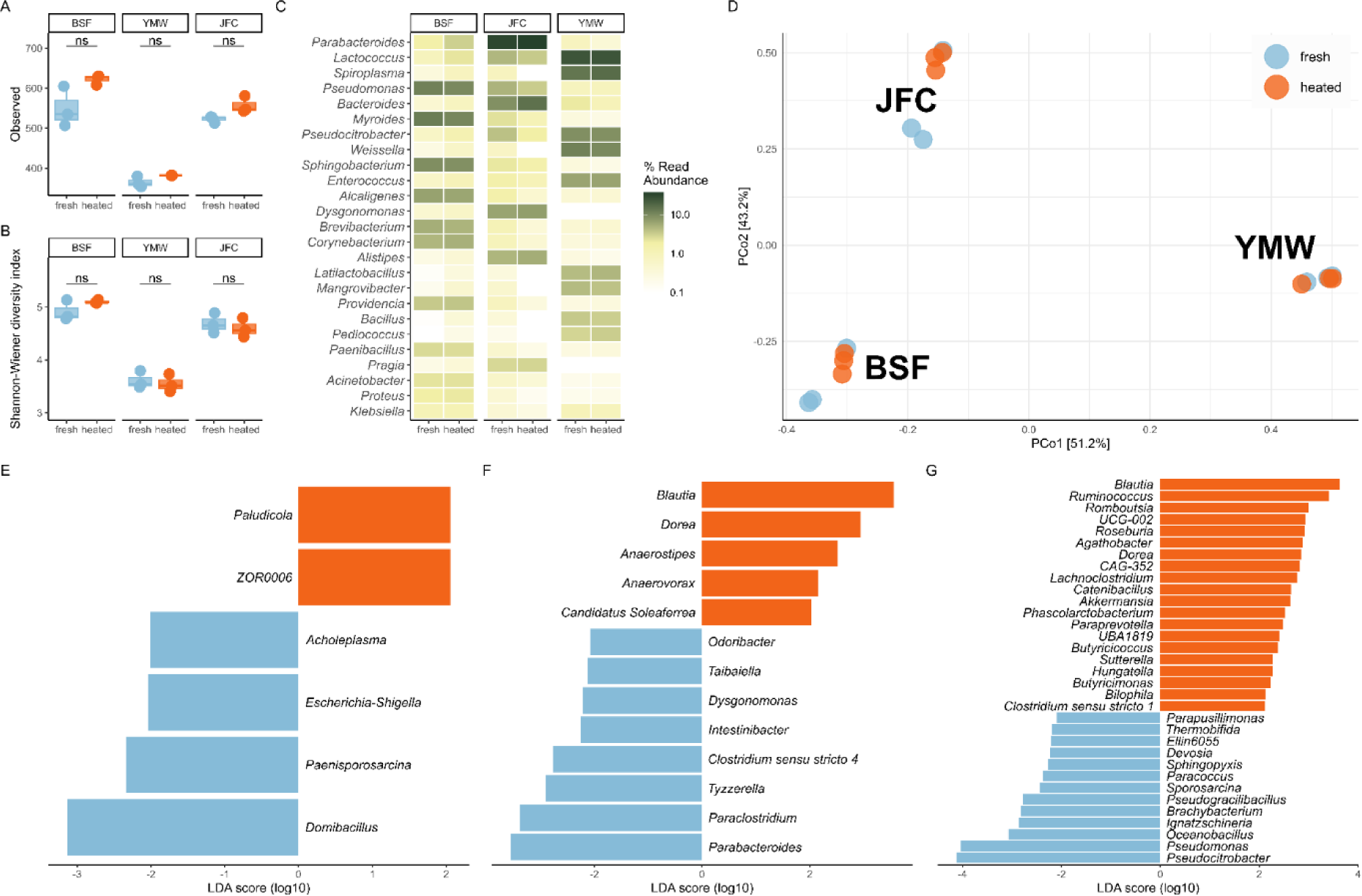
Analysis of microbial communities in fresh (blue) and heat-treated (orange) frass samples of black soldier fly (BSF), yellow mealworm (YMW), and Jamaican field cricket (JFC) (n = 3). Alpha diversity explained by (**A**) observed species and (**B**) Shannon-Wiener diversity index. **C**. The 25 most abundant genera based on relative abundance. ASVs without classification to the genus level were removed from the list. **D**. Principal coordinates analysis (PCoA) showing the species- and treatment-derived (dis)similarity between samples based on Bray-Curtis dissimilarity. Differentially abundant bacterial genera in BSF (**E**), YMW (**F**), and JFC (**G**) samples as determined via linear discriminant analysis of effect size (LEfSe).

The differences in the frass’ microbiome composition among the three insect species were further illustrated by the diverging insect species-specific patterns in the relative abundances of the top 25 bacterial genera (Figure C) and the distance between sample aggregates visualized by the PCoA (Figure 4D). PERMANOVA validated the presence of significant differences among the frass microbiomes of the three insect species, both in their fresh state (*F_2,6_* = 51.82, p < 0.01) or after heat treatment (*F_2,6_* = 51.82, p < 0.01). The most dominant genera for BSF, JFC, and YWM frass were *Pseudomonas*, *Parabacteroides*, and *Lactococcus*, respectively.

Little overlaps were found in biomarker genera characterizing fresh and heat-treated samples of BSF (Figure 4E), YMW (Figure 4F), and JFC (Figure 4G). Most differentially abundant genera explaining the divergence in microbiome composition between fresh and heated frass were found in JFC samples. Only two genera (*Blautia* sp. and *Dorea* sp.) characteristic for heat-treated JFC frass were also found in heat-treated YMW samples. The least characteristic genera were found for BSF samples. Bacterial genera explaining the overall differences in frass microbiomes among the three insect species are reported in Supplementary Figure 2.

### 3.4. Physicochemical characterization of frass-supplemented soils and control soils

To assess the effect of frass supplementation on the soil, the soil:frass mixtures were analyzed after a two-week greenhouse incubation period and compared both to each other and to the control soil. Throughout the incubation, the soil moisture within the pots decreased by max. 25%. To maintain consistency, the soil moisture was continuously adjusted back to its initial value (Supplementary Figure 1). Consequently, at the end of the incubation period, the water content and TS in the frass-treated soil samples were comparable to the control soil (Table 5).

**Table 5.** Physicochemical parameters (mean ± standard deviation) measured in the control soil and the soil:frass mixtures after two-week incubation in a greenhouse at 20 °C. Statistical differences were calculated via analysis of variance (ANOVA) followed by Tukey HSD posthoc tests for pairwise comparison of sample groups (n = 4). b.d.l. = below detection limit.

|  | Control soil | Soil:Frass (BSF) |  | Soil:Frass (YMW) |  | Soil:Frass (JFC) |  | *F*-value |
| --- | --- | --- | --- | --- | --- | --- | --- | --- |
|  |  | Fresh | Heated | Fresh | Heated | Fresh | Heated |  |
| Water content [%] | 24.8 ± 0.4<sup>c</sup> | 25.0 ± 0.4<sup>bc</sup> | 24.3 ± 0.2<sup>c</sup> | 26.2 ± 0.4<sup>a</sup> | 26.1 ± 0.4<sup>a</sup> | 26.1 ± 0.8<sup>ab</sup> | 24.2 ± 0.4<sup>c</sup> | *F*=15.33*** |
| Total solids [%] | 75.24 ± 0.35<sup>a</sup> | 74.97 ± 0.44<sup>ab</sup> | 75.73 ± 0.21<sup>a</sup> | 73.8 ± 0.4<sup>c</sup> | 73.9 ± 0.4<sup>c</sup> | 73.9 ± 0.8<sup>bc</sup> | 75.8 ± 0.4<sup>a</sup> | *F*=15.33*** |
| Volatile solids [%] | 6.23 ± 0.13<sup>e</sup> | 7.14 ± 0.32<sup>d</sup> | 7.07 ± 0.15<sup>d</sup> | 11.09 ± 0.25<sup>a</sup> | 10.97 ± 0.49<sup>ab</sup> | 10.21 ± 0.36<sup>b</sup> | 9.19 ± 0.56<sup>c</sup> | *F*=128.081*** |
| Ash [%] | 69.0 ± 0.4<sup>a</sup> | 67.8 ± 0.5<sup>ab</sup> | 68.7 ± 0.4<sup>a</sup> | 62.7 ± 0.3<sup>c</sup> | 62.9 ± 0.8<sup>c</sup> | 63.7 ± 1.0<sup>c</sup> | 66.6 ± 0.5<sup>b</sup> | *F*=88.858*** |
| C [%] | 4.80 ± 0.27<sup>b</sup> | 5.55 ± 0.29<sup>b</sup> | 5.51 ± 0.16<sup>b</sup> | 7.59 ± 0.54<sup>a</sup> | 7.43 ± 0.3<sup>a</sup> | 7.19 ± 0.23<sup>a</sup> | 7.46 ± 0.63<sup>a</sup> | *F*=37.84*** |
| N [%] | 0.31 ± 0.02<sup>b</sup> | 0.38 ± 0.01<sup>b</sup> | 0.38 ± 0.01<sup>b</sup> | 0.59 ± 0.04<sup>a</sup> | 0.59 ± 0.02<sup>a</sup> | 0.61 ± 0.04<sup>a</sup> | 0.63 ± 0.05<sup>a</sup> | *F*=69.761*** |
| C:N ratio | 15.5 ± 2.0<sup>a</sup> | 14.6 ± 0.8<sup>ab</sup> | 14.5 ± 0.7<sup>ab</sup> | 12.9 ± 0.8<sup>bc</sup> | 12.5 ± 0.2<sup>bc</sup> | 11.8 ± 1.0<sup>c</sup> | 11.9 ± 0.6<sup>c</sup> | *F*=8.585*** |
| NH<sub>4</sub>-N [μg g<sup>-1</sup> TS] | 1.9 ± 0.4<sup>c</sup> | 12.4 ± 1.8<sup>c</sup> | 12.6 ± 0.6<sup>c</sup> | 821.3 ± 41.8<sup>b</sup> | 788.6 ± 41.4<sup>b</sup> | 1690.5 ± 37.4<sup>a</sup> | 1757.3 ± 73.2<sup>a</sup> | *F*=1619.117*** |
| NO<sub>2</sub>-N + NO<sub>3</sub>-N [μg g<sup>-1</sup> TS] | 34.1 ± 4.5<sup>c</sup> | 243.3 ± 38.1<sup>b</sup> | 231.8 ± 15.7<sup>b</sup> | 386.7 ± 15.7<sup>a</sup> | 377.2 ± 41.2<sup>a</sup> | 8.9 ± 0.8<sup>c</sup> | 4.9 ± 1.2<sup>c</sup> | *F*=215.528*** |
| P<sub>plant available</sub> [μg g<sup>-1</sup> TS] | 9.0 ± 0.2<sup>c</sup> | 72.1 ± 21.3<sup>c</sup> | 64.7 ± 9.9<sup>c</sup> | 788.6 ± 66.5<sup>a</sup> | 861.6 ± 40.1<sup>a</sup> | 583.2 ± 49.0<sup>b</sup> | 613.5 ± 69.1<sup>b</sup> | *F*=275.029*** |
| pH | 8.06 ± 0.05<sup>c</sup> | 7.62 ± 0.03<sup>d</sup> | 7.56 ± 0.01<sup>d</sup> | 8.18 ± 0.1<sup>bc</sup> | 8.26 ± 0.03<sup>b</sup> | 8.59 ± 0.08<sup>a</sup> | 8.54 ± 0.05<sup>a</sup> | *F*=189.352*** |
| EC [μS cm<sup>-1</sup>] | 84.1 ± 4.2<sup>d</sup> | 225.3 ± 36.3<sup>c</sup> | 222.5 ± 10.3<sup>cd</sup> | 882.3 ± 45.1<sup>b</sup> | 820.8 ± 50.6<sup>b</sup> | 1099.3 ± 113.5<sup>a</sup> | 1201.0 ± 84.7<sup>a</sup> | *F*=227.595*** |
| Salinity | b.d.l.<sup>c</sup> | b.d.l.<sup>c</sup> | b.d.l.<sup>c</sup> | 0.2 ± 0<sup>b</sup> | 0.18 ± 0.05<sup>b</sup> | 0.33 ± 0.05<sup>a</sup> | 0.38 ± 0.05<sup>a</sup> | *F*=94.444*** |
| TDS [mg L<sup>-1</sup>] | 33.8 ± 1.7<sup>c</sup> | 89.8 ± 14.2<sup>c</sup> | 89.0 ± 4.6<sup>c</sup> | 353.0 ± 18.1<sup>b</sup> | 328.3 ± 19.9<sup>b</sup> | 440.3 ± 45.5<sup>a</sup> | 480.0 ± 33.7<sup>a</sup> | *F*=228.644*** |

With around 6%, the VS content in the control soil was ca. 1% lower than in soils treated with BSF frass and 3-5% lower than in soils treated with JFC and YMW frass, respectively. The supplementation of YMW and JFC frass resulted in a significant increase in the soil’s C and N content by up to 2.8% and 0.3%, respectively, leading to a decrease in the C:N ratio from 15.5 to a minimum of 11.8 in soil treated with fresh JFC frass.

Although NH_4_^+^-N concentrations were initially highest in samples of fresh and heated BSF frass (Table 3), they exhibited the lowest levels in soils supplemented with BSF frass, given the recommended dosage used. The JFC frass application (12.5% fresh or 11.6% heated, w/w) resulted in the highest soil NH_4_^+^-N concentration of 1723.9 µg g^-1^ TS and, thus, to an increase of 900% of the original soil concentration after two weeks and a one-time amendment. YMW frass, having the highest NO_2_-NO_3_--N content, increased the soil NO_2_-NO_3_--N levels by ca. 350 µg g^-1^ TS compared to control soil. Plant-available P reached on average 11 mg g^-1^ TS (YMW, JFC) and 20 mg g^-1^ TS in raw BSF frass. Taking the concentration of the amended 2% (BSF), an average of 12% (YMW and JFC) and the original soil concentration into account, resulting plant-available P concentrations ranged from 15, 64 to 38% of the initially applied P in BSF, YMW, and JFC treatments, respectively. Thus, significantly higher levels of available P in soils supplemented with YMW frass (789 and 862 µg P g^-1^ TS for fresh and heated frass, respectively) were established compared to soils supplemented with BSF (72 and 65 µg P g^-1^ TS for fresh and heated frass, respectively) and JFC frass (583 and 614 µg P g^-1^ TS for fresh and heated frass, respectively). The ratios of plant-available N (NH_4_^+^-N and NO_2_-NO_3_--N) to P (N:P) in the frass-amended soils reached 1.4 (YMW), 2.9 (JFC), and 3.6 (BSF). While YMW and JFC frass led to a slight increase in soil pH to 8.6 after treatment, the addition of BSF frass caused a slight decrease in pH from 8.0 to 7.5. EC and TDS increased in all frass-treated soils, with maxima of 1201 µS cm^-1^ and 480 mg L^-1^ measured after the supplementation of heat-treated JFC frass.

### 3.5. Microbiological characterization of frass-supplemented soils and control soils

In contrast to the microbial activity observed in fresh and heat-treated BSF frass, its application to the soil did not lead to significant promotion of BR and C_mic_. As a result, the MQ in the treated soil was slightly lower compared to the control (Table 6). However, the supplementation of YMW and JFC frass led to a significantly higher soil microbial activity. While BR was comparable in soils supplemented with both fresh and heat-treated YMW and JFC frass, the C_mic_ was remarkably higher in soils mixed with fresh and heat-treated YMW frass. The comparable BR rates but lower C_mic_ in soils containing JFC frass, in turn, resulted in higher MQ values.

**Table 6.**
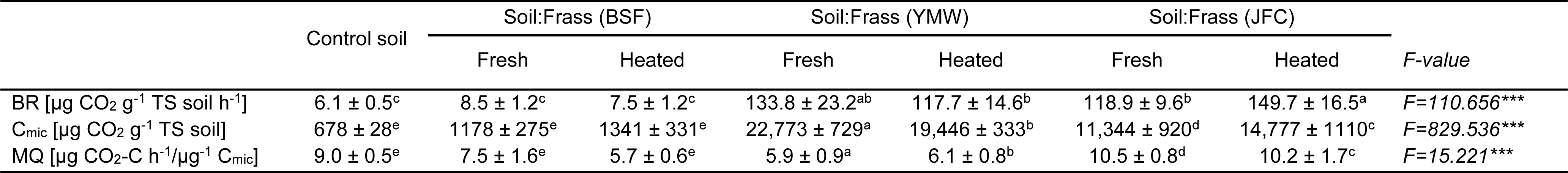
Microbiological parameters (mean ± standard deviation) before and after treating the frass at 70 °C for 1 hour. Statistical differences were calculated via analysis of variance (ANOVA) followed by Tukey HSD posthoc tests for pairwise comparison of sample groups (n = 4).

Although no colonies identified as *E. coli* were detected in any of the raw frass samples, abundances of 1.5 10^4^ and 9.8 10^3^ CFUs g^-1^ TS soil of *E. coli* were found in soils mixed with fresh BSF and YMW frass, respectively (Figure 3). Neither the control soil nor the soils supplemented with JFC frass showed any growth of *E. coli*. Initially, *Salmonella* sp. was exclusively detected in fresh JFC frass and not in any other frass samples. However, following the two-week soil incubation period, *Salmonella* sp. was found in soils mixed with fresh BSF, fresh or heat-treated YMW, and heat-treated JFC in numbers ranging between 5.0 10^2^ to 2.2 10^4^ CFUs g^-1^ soil_TS_. No *Salmonella* sp. was detected in the control soil.

## 4. Discussion

### 4.1. Frass exhibits multi-level variations depending on the insect species

The physicochemical and microbiological characterization of black soldier fly (BSF), yellow meal worm (YMW), and Jamaican field cricket (JFC) frass has shown that the frass’ properties are intrinsically related to the insect species (Figure 2 and Table 3). This goes along with prior studies showing that the differences in feed requirements and rearing conditions are reflected in the properties of the frass (Beesigamukama et al., 2022). At the physicochemical level, significantly higher moisture content in BSF frass can be explained by the faster development of the larvae compared to YMW and JFC which typically take three times longer before reaching a harvest-ready stage, thus leaving less time for the substrate to dry. The frass moisture content at the time of harvest emerges as a crucial parameter that influences the separability of insects and rearing residues by sieving (Gärttling and Schulz, 2022). Operators face the challenge of finely adjusting the initial substrate moisture to establish suitable conditions throughout the rearing process while at the same time considering water loss through evaporation, uptake from insects, and metabolization by microorganisms, as this affects the successive processing steps.

At the microbiological level, fresh BSF frass stood out with significantly higher microbial activity along with CFU counts of viable aerobic microorganisms that were up to four orders of magnitudes higher than in fresh YMW and JFC frass. This disparity in microbial growth and metabolism is likely sustained by the higher moisture content in BSF frass.

At the microbiome level, dominant genera did not overlap among insect species, underscoring that different types of frass exhibit unique microbial signatures (Figure 4C).

### 4.2. Heat treatment has limited impact on frass nutrients but reduces microbial activity and viable counts of pathogenic microbes

The primary rationale for heat-treating insect frass is to guarantee its safety by removing any potential microbial pathogens present within these residues. Notably, in the EU, frass has recently been classified in the same category as processed manure, thus requiring further treatment (Regulation (EU) 2021/1925, 2021). The selection of substrates authorized for insect rearing remains considerably constrained when compared with the extensive range of organic wastes deemed suitable for this purpose, but microbes residing in insect guts are inevitably transferred into the frass via excretion. However, the prescribed heat treatment (70 °C for 1 h) seems to be appropriate to reduce the microbial load beneath the detection limit of all tested frass types, which cover a broad range of species that are currently mass-reared in Europe for feed and food purposes.

While heat treatment may be efficient in hygienizing frass from a microbiological perspective, the effect of temperature on other potential pollutants in frass should be considered. Organic wastes suitable as rearing substrate are prone to be contaminated with microplastics. While first studies suggest that the development of farmed insects is not significantly affected by microplastics, these particles are excreted in their original form after passing through the larvae’s digestive tract and accumulate in the frass (Heussler et al., 2023; Lievens et al., 2023). Temperatures exceeding 60 °C have been shown to melt or clump specific microplastics, and temperatures nearing 100 °C might even lead to their elimination (Munno et al., 2018). Nonetheless, the potential impacts on frass production are yet to be explored.

### 4.3. Frass supplementation improves plant nutrient content and microbial activity in soils

Frass, as the main byproduct of insect rearing processes, has the potential to be used as a soil improver and plant fertilizer, by supplying soil particularly with nitrogen (N), phosphorus (P) and potassium (K) (Fuertes-Mendizábal et al., 2023; Klammsteiner et al., 2020a). The recent EU regulation (Regulation (EU) 2021/1925, 2021) foresees the heat treatment of frass to guarantee safety upon consumption of crops and plants fertilized by frass. Nutrient conditions in soils amended with fresh and heat-treated frass were significantly improved in both cases and heat treatment did not significantly alter the improvement, irrespective of frass type. According to our knowledge, this is the first report of amended frass after heat treatment to soil, demonstrating that the fertilizer capacity of frass stays unaltered after one-time heating.

Generally, the nutrient load of the frass is comparable to other organic fertilizers, like compost (Poletschny, 1994). Thus, fertilization with frass exhibits highly favorable attributes with regard to all essential nutrients, including carbon (C), N, and P. In terms of C and N content, frass aligns with concentrations established in other organic fertilizers (Poletschny, 1994). The utilization of frass is scalable, and like other organic fertilizers, it has a sustainable impact on soil in contrast to mineral fertilizers. This sustainability arises from the fact that nutrients are mobilized by microorganisms and/or are incorporated into microbial biomass. This was particularly pronounced in the case of the N fractions, where the average amendment of 12% (w/w) of YMW and JFC frass to soil led to a strong mineralization, resulting in the transfer of a relevant fraction of total N into NH_4_^+^-N. Consequently, the soil:frass mixture contained up to six times more NH_4_^+^-N than the initial addition would correspond to. Similarly important for plant nutrition, albeit less pronounced here, this was observed for available P as well. Particularly promising is the high amount of plant-available P of all three frass types as, after nitrogen, P is the second most limited nutrient and is not available for the plant despite abundant phosphorus reserves (Illmer and Schinner, 1992).

In soil, phosphate is usually present as insoluble aluminum-, iron- and calcium phosphate (Kooijman et al., 2002). Due to its insoluble form, P fertilizers are commonly used in agriculture to increase crop productivity (Ros et al., 2020). In fact, the available P content in the frass exceeded concentrations by approximately 4-fold when compared to typical total P levels found in organic fertilizers. This high amount of P is also reflected in the available P that was present in the soil after fertilization. Two weeks after a single application, P levels reached up to 860 µg g^-1^ TS weight, increasing soil phosphorus by 100-1000 times compared to the current control and available P in soils globally, respectively (McDowell et al., 2023). The P fertilization associated with frass is consistently beneficial in all cases and supports the replacement or reduction of chemical P fertilizers which is a major goal in sustainable agriculture. Furthermore, the increased concentration of phosphorus established in the frass amended soils could affect soil N pools and processes positively as well (Wang et al., 2022). Still, consideration should be given to whether the elevated P levels are ecologically feasible.

Besides the added nutrients during the single frass application, the supplementation of frass boosted soil microbial activity especially for YMW and JFC treatments. The lower performance of BSF frass can be associated with the lower dosage recommended by the producers. The recommended high-dosage applications of YMW and JFC refer to small-scale garden practices, thus, large-scale applications will be lower as a linear up-scaling of the necessary amount of frass will lead to unfeasible amounts. The nutrient support by frass amendment and the autochthonous microbes in the frass significantly increased the microbial biomass in the soils in case of JFC and YMW frass. Despite the absence of detectable physiologically active microbial biomass in the case of raw YMW and JFC frass, the addition to the soil significantly boosted microbial activity and biomass which can be traced back to the associated increase in water availability for the frass microbes and the high nutrient input, indicating that not only did the frass provide nutrients, but it also benefited the soil’s resident microbes. The heat-treatment led to contrasting and frass-type specific effects, still providing very similar positive effects on soil quality compared to untreated fresh frass.

## 5. Conclusion

Variations among frass samples were primarily attributed to the insect species, with minimal influence from heat treatment. While BSF frass demonstrated the highest NH_4_^+^-N concentrations, levels of NO_2_-NO_3_--N were significantly elevated in YMW frass. Interestingly, BSF frass also exhibited the highest content of plant-available P, even after heat-treatment, and it was the only frass type displaying microbial activity that significantly decreased following heat treatment. Supplementing soil with frass led to distinct shifts in soil properties, with YMW frass having the most wide ranging effects on nutrient concentrations. Collectively, these findings provide substantial insights into the intricate interactions between insect frass, heat treatment, and soil dynamics, with potential implications for sustainable agricultural practices.

## 6. Data availability statement

Raw NovaSeq6000 amplicon data were deposited into the European Nucleotide Archive (ENA) under the accession number PRJEB61412 and are available at the following URL: https://www.ebi.ac.uk/ena/browser/view/PRJEB61412.

## 9. Author contributions

**Nadine Praeg**: Conceptualization, Methodology, Validation, Formal analysis, Investigation, Resources, Writing - Original Draft, Writing - Review & Editing. **Thomas Klammsteiner**: Conceptualization, Methodology, Validation, Formal analysis, Investigation, Resources, Data Curation, Writing - Original Draft, Writing - Review & Editing, Visualization, Supervision, Project administration, Funding acquisition.

## Acknowledgements

We would like to express our gratitude to Maria Payr and Max Moser for their helpful assistance in the laboratory, and to Paul Illmer for his advice in collecting the soil for this study and data interpretation. We also thank Aleksandar Gavrilović and Simon Weinberger/Ecofly GmbH for generously providing us with fresh insect frass.

