## Supplemental Figures and Tables for "Frass fertilizers from mass-reared insects: species variation, heat treatment effects, and implications for soil application"

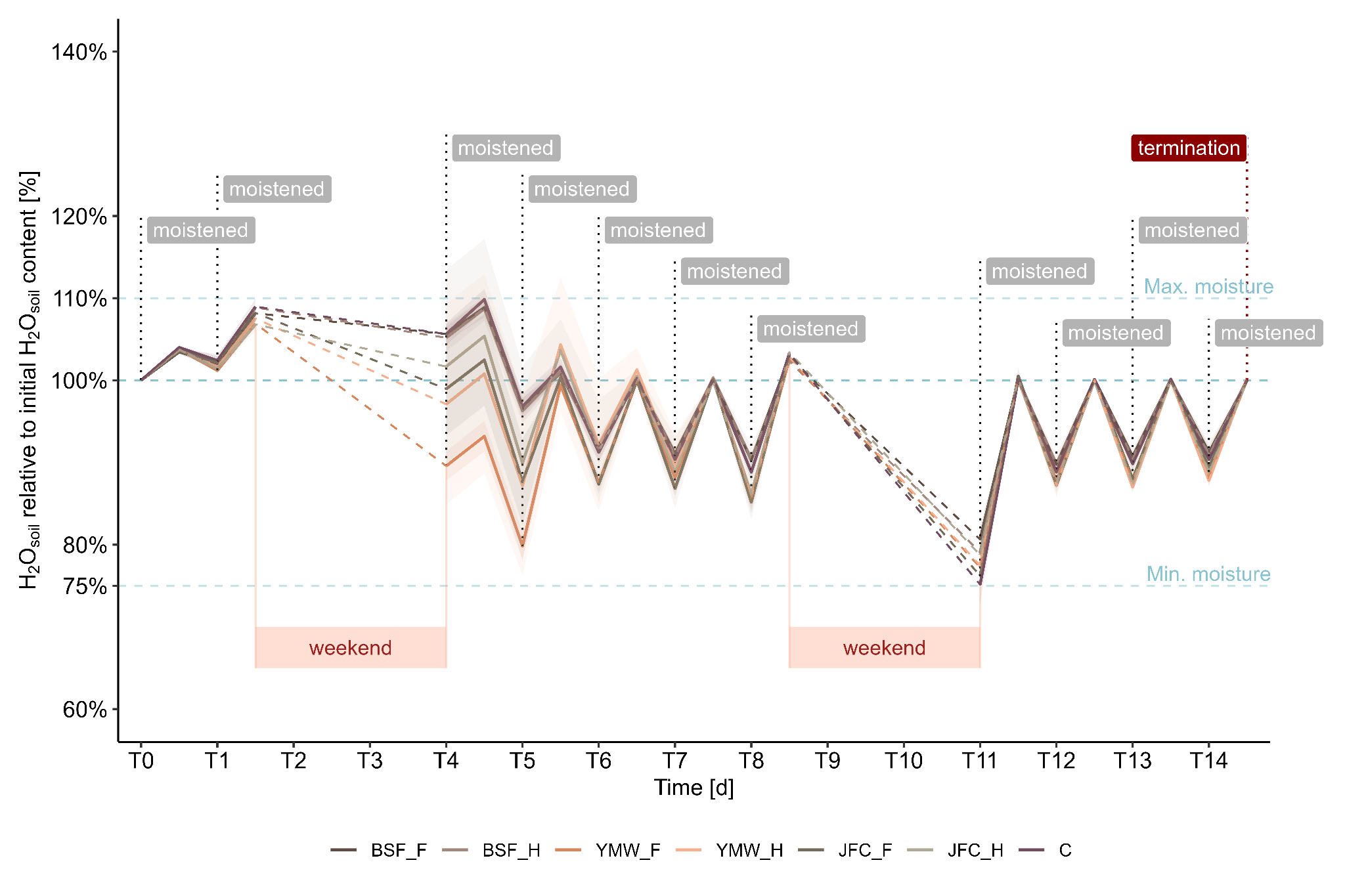

**Supplementary figure 1**. During the 14-day greenhouse incubation trial, soil moisture was monitored. Daily measurements of water loss through evaporation were taken on workdays and used to restore the soil water content to its original level by spraying the surface with distilled water. BSF_F and BSF_H = soil with fresh and heat-treated black soldier fly frass, respectively; YMW_F & YMW_H = soil with fresh and heat-treated yellow mealworm frass, respectively; JFC_F and JFC_H = soil with fresh and heat-treated Jamaican field cricket frass, respectively; C = control soil.

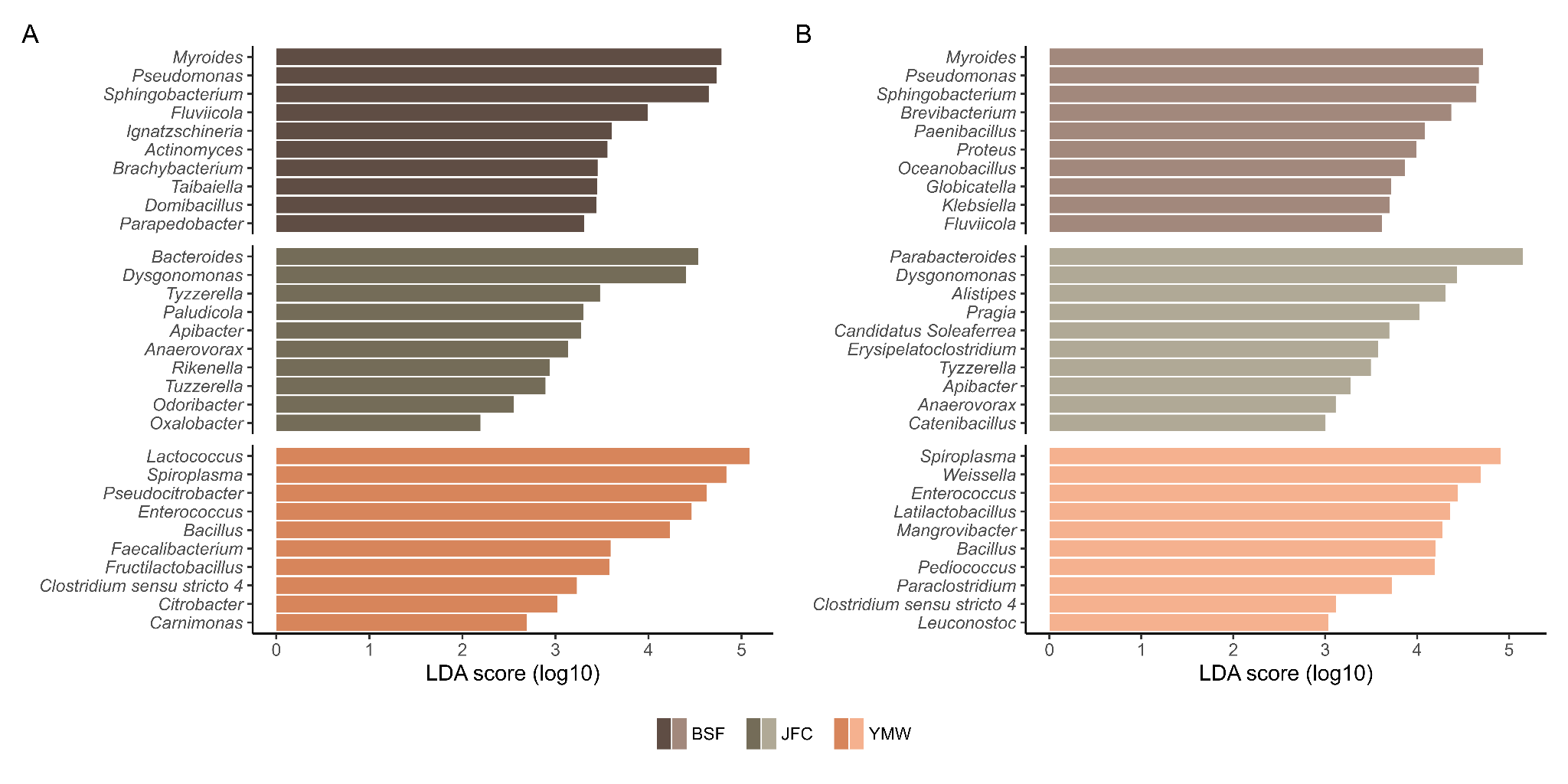

**Supplementary figure 2.** Top 10 differentially abundant genera identified by linear discriminant analysis of effect size (LEfSe) in fresh (**A**) and heat-treated (**B**) frass samples of black soldier fly (BSF), yellow mealworm (YMW), and Jamaican field cricket (JFC).

**Supplementary Table 1.** Average counts (mean ± standard deviation) of colony forming units (CFUs) per g TS of frass after plating samples on standard (STD) I, ChromoCult™ TBX, and XLT4 agar and incubating for 48 h. Statistical differences were calculated via analysis of variance followed by Tukey HSD posthoc tests for pairwise comparison of sample groups (n = 3). b.d.l. = below detection limit.

| Medium | BSF frass | | YMW frass | | JFC frass | | *F-value* |
| --- | --- | --- | --- | --- | --- | --- | --- |
|  | Fresh | Heated | Fresh | Heated | Fresh | Heated |  |
| STD I agar CFU | 1.3 10^9^ ± 5.8 10^7a^ | 3.8 10^8^ ± 1.5 10^8b^ | 8.2 10^5^ ± 9.0 10^4c^ | 4.9 10^5^ ± 1.3 10^5c^ | 5.7 10^5^ ± 1.3 10^5c^ | 2.6 10^5^ ± 6.5 10^4c^ | *F=172.13**** |
| TBX CFU | 4.8 10^6^ ± 1.3 10^6a^ | 1.6 10^6^ ± 6.9 10^5b^ | 1.6 10^5^ ± 4.4 10^4b^ | 8.3 10^4^ ± 2.2 10^4b^ | 3.0 10^5^ ± 1.7 10^5b^ | 2.5 10^4^ ± 2.8 10^4b^ | *F=27.271**** |
| *E. coli* | b.d.l. | b.d.l. | b.d.l. | b.d.l. | b.d.l. | b.d.l. |  |
| XLT4 CFU | 2.8 10^5^ ± 7.6 10^4a^ | 4.0 10^5^ ± 8.3 10^4a^ | 4.9 10^4^ ± 1.2 10^4b^ | 1.7 10^4^ ± 1.8 10^3b^ | 6.3 10^4^ ± 1.1 10^4b^ | 1.7 10^4^ ± 2.8 10^3b^ | *F=36.697**** |
| *Salmonella* sp. | b.d.l. | b.d.l. | b.d.l. | b.d.l. | 1.7 10^3^ ± 1.7 10^3^ | b.d.l. |  |

**Supplementary Table 2.** Average counts (mean ± standard deviation) of colony forming units (CFUs) per g TS of frass after plating samples on ChromoCult™ TBX and XLT4 agar and incubating for 48 h. Statistical differences were calculated via analysis of variance (ANOVA) followed by Tukey HSD posthoc tests for pairwise comparison of sample groups (n = 3). n.c. = not countable, b.d.l. = below detection limit.

| Medium | Control soil | Soil:Frass (BSF) | | | Soil:Frass (YMW) | | Soil:Frass (JFC) | | *F-value* |
| --- | --- | --- | --- | --- | --- | --- | --- | --- | --- |
|  |  | Fresh | Heated | Fresh | | Heated | Fresh | Heated |  |
| TBX CFU | 8.5 10^4^ ± 1.9 10^4^ | n.c. | n.c. | n.c. | | n.c. | n.c. | n.c. |  |
| *E. coli* | b.d.l.^a^ | 1.5 10^4^ ± 2.6 10^4a^ | b.d.l.^a^ | 9.8 10^3^ ± 8.5 10^3a^ | | b.d.l.^a^ | b.d.l.^a^ | b.d.l.^a^ | *F=1.09* |
| XLT4 CFU | 3.5 10^4^ ± 2.1 10^4c^ | 1.3 10^5^ ± 6.3 10^4c^ | 2.3 10^5^ ± 5.5 10^4c^ | 1.3 10^6^ ± 3.1 10^5a^ | | 7.8 10^5^ ± 3.3 10^5b^ | 4.2 10^5^ ± 1.4 10^5bc^ | 4.2 10^5^ ± 6.4 10^4bc^ | *F=17.665**** |
| *Salmonella* sp. | b.d.l.^a^ | 5.0 10^2^ ± 8.7 10^2a^ | b.d.l.^a^ | 7.8 10^3^ ± 1.1 10^4a^ | | 2.2 10^4^ ± 3.3 10^4a^ | b.d.l.^a^ | 2.0 10^3^ ± 3.5 10^3a^ | *F=1.143* |
